# An efficient beet severe curly top virus-based VIGS vector in *Beta vulgaris*

**DOI:** 10.1101/2025.07.11.663803

**Authors:** Zhangyao Nie, Zhihong Guo, Xinyu Qin, Qi Liu, Minglongcheng Zhou, Jian Ye, Zongying Zhang, Chenggui Han, Ying Wang

**Affiliations:** Ministry of Agriculture and Rural Affairs Key Laboratory of Pest Monitoring and Green Management, College of Plant Protection, China Agricultural University, Beijing, China; State Key laboratory of Plant Genomics, Institute of Microbiology, Chinese Academy of Sciences, Beijing, China

**Keywords:** sugar beet, BSCTV, VIGS vector, RNA silencing

## Abstract

Sugar beet (*Beta vulgaris* L.) is one of the most important sugar crops worldwide. However, studies on sugar beet gene functions are still lagging compared to other crops due to the lack of an effective genetic manipulation. In this study, we generated an infectious clone of beet sever curly top virus (BSCTV) and engineered it into a VIGS vector. BSCTV-*BvPDS* can induce strong and persistent bleached *PDS*-silencing phenotype in 8 tested sugar beet varieties. We further efficiently silenced *B. vulgaris MYB1* gene that regulating the betalain pathway in sugar beet by BSCTV-based VIGS system. Finally, we used this system to silence the effector *CbNip1* of *Cercospora beticola* through cross-kingdom RNAi, resulting in reduced disease symptoms in sugar beet plants. Therefore, the BSCTV VIGS system can be used to study sugar beet-pathogen interactions. In summary, we have established an easy and efficiency BSCTV-based VIGS vector in sugar beet. This system will be an attractive tool for functional genomic studies for sugar beet.

## Brief Communication

Sugar beet (*Beta vulgaris* L.), one of the most important sugar crops worldwide, provides 20-30% of the global sugar supplies (FAO, 2023). However, genomic studies on sugar beets are very limited due to the lack of efficient genetic systems. Virus-induced gene silencing (VIGS) has been widely exploited to investigate gene function in various plant species (Liu *et al*., 2002). Although VIGS is applicable in inoculated leaves of sugar beet (Hatlestad *et al*., 2015), virus-induced highly efficient and systemic RNA silencing is urgently needed. Beet severe curly top virus (BSCTV) is a monopartite virus belonging to the genus *Curtovirus* (family *Geminiviridae*). BSCTV has a very wide host range within 44 dicot plant families (Strausbaugh *et al*., 2008). Thus, BSCTV is a good candidate virus to be developed into VIGS vectors for studying sugar beet and other plant species.

In this study, we first inserted 1.2 copies of BSCTV genome (GenBank accession: U02311) into a binary vector pCB301 to generate an infectious cDNA clone (pCB-BSCTV, Figure 1a). The bacterium containing pCB-BSCTV was infiltrated into leaves of sugar beet variety BETA176. At 12dpi, the rescued BSCTV induced severe leaf curling symptoms (Figure 1b) and exhibited high infectivity in the systemic leaves and roots of sugar beet (Figure S1a). Since BSCTV C2 protein is a determiner of viral pathogenicity and the C1/C2/C3 overlapping region also produces abundant of viral small interfering RNAs (vsiRNAs) (Zhang *et al*., 2011; Voorburg *et al*., 2021), the overlapping region could be used as the insertion site to decrease viral symptom and induce strong RNA silencing. To this end, the 300 bp GFP and *B. vulgaris Phytoene desaturase* (*BvPDS*) fragments were independently inserted at C1/C3 junction in pCB-BSCTV (Figure 1a). As expected, BSCTV-*GFP* and BSCTV-*BvPDS* induced mild symptoms through disrupting the C2 coding frame (Figure 1c). Furthermore, systemically infected leaves of sugar beet inoculated with BSCTV-*BvPDS* exhibited bleaching phenotype at 20 dpi (Figure 1c). Subsequently, the bleached phenotype was extended to all new emerging infected leaves (Figure 1d). RT-qPCR results revealed a significantly down-regulated level of *BvPDS* in BSCTV-*BvPDS* infected plants compared that of BSCTV-*GFP*-treated plants (Figure 1e). The inserted fragments of *GFP* and *BvPDS* were genetically stable at 30 dpi (Figure 1f). Moreover, the results of four independent experiments showed that ∼ 82% sugar beet plants (n = 55) exhibited bleached phenotype after induction by BSCTV-VIGS (Table S1). We also confirmed that 180 bp of *BvPDS* fragment was sufficient to induce target gene silencing (Figure S1b). Next, BSCTV-*BvPDS* could apply to 8 tested sugar beet varieties exhibiting bleached phenotypes (Figure 1g). Strikingly, SV1434 and table beet showed fully bleached phenotypes (Figure 1g). These results demonstrate that BSCTV-VIGS can be engineered for genomic studies of sugar beet.

**Figure 1.**
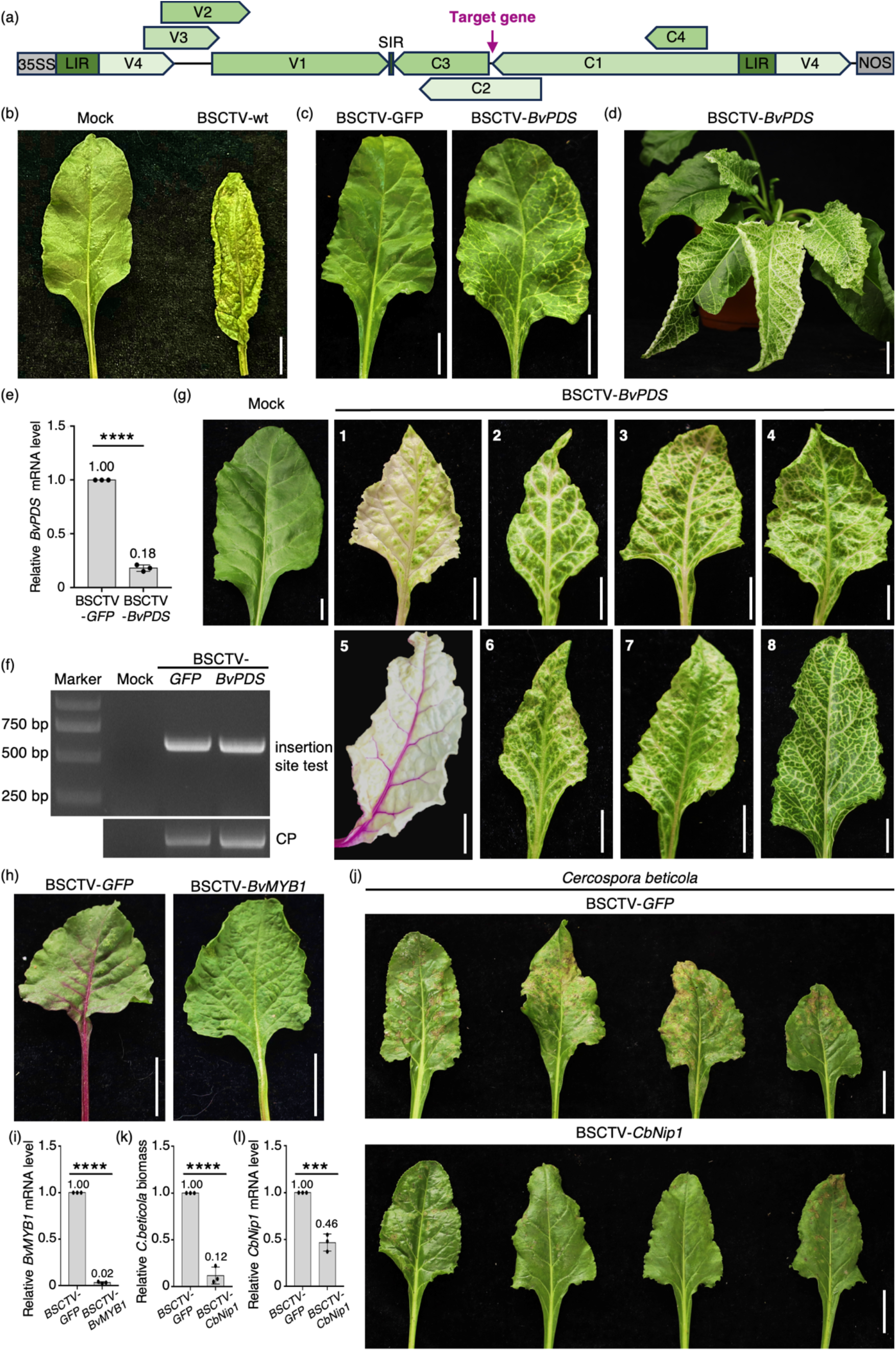
The BSCTV-based VIGS system in sugar beet. (a) Schematic diagram showing the construction strategy of the full-length BSCTV infectious cDNA clone. The arrow indicates the insertion site. (b) Systemic symptoms in sugar beet plants after agroinfiltration of BSCTV-wt. Scale bars, 1 cm (c) The bleached phenotype was first observed on sugar beet leaves 20 days after agroinfiltration with BSCTV-*BvPDS*. Scale bars, 2 cm. (d) The silencing symptoms on the sugar beet plants after 6 weeks of agroinfiltration with BSCTV-*BvPDS*. Scale bars, 2.5 cm (e) The RT-qPCR analysis confirmed *BvPDS* was silenced after the agroinfiltration of BSCTV-*BvPDS*. The data were analysed using the Student’s t-test (two-tailed, *P < 0.05, ***P < 0.001, ****P < 0.0001). (f) The PCR detection indicated that the BSCTV VIGS vector did not revert to the wild type. (g) The bleaching symptoms on different varieties of sugar beet. 1: SV1434; 2: KWS-2407; 3: KVHN-4092; 4: 94004-2. 5: table beet; 6: KWS-7747; 7: MA-3028; 8: BETA176. Scale bars, 1 cm. (h) The *BvMYB1* silencing symptoms on the leaf of table beet. Scale bars, 2 cm. (i) The RT-qPCR analysis confirmed *BvMYB1* was silenced in the leaves of table beet after agroinfiltration with BSCTV-*BvMYB1*. (j) The infection symptoms on the sugar beet leaves, 10 days after inoculation with *Cercospora beticola*. Scale bars, 2.5 cm. (k, l) The qPCR analyses confirmed that the treatment with agroinfiltration of BSCTV-*CbNip1* resulted in a lower relative biomass of *C. beticola*, and the target gene *CbNip1* was silenced.

We further tested BSCTV-based vector for other genes. We firstly silenced *B. vulgaris MYB1* (*BvMYB1*) that regulating the betalain pathway in sugar beets (Hatlestad *et al*., 2015). Therefore, a 300 bp *BvMYB1* fragment was inserted into pCB-BSCTV that was agroinfiltrated into sugar beets. The plants infected with BSCTV-*BvMYB1* displayed green veins and stems of systemic leaves compared with BSCTV-*GFP* infected plants at 35 dpi (Figure 1h). RT-qPCR results confirmed a significant down-regulation of *BvMYB1* accumulation in leaf tissues compared to those of BSCTV-*GFP* infected plants (Figure 1i). These results demonstrate that the BSCTV-based VIGS system can be applied to gene function studies in leaf and stem of sugar beet plants.

We next used BSCTV VIGS vector to cross-kingdom silence vital genes of *Cercospora beticola*, a fungus that causes the most destructive foliar disease of sugar beet. We engineered the recombinant vector to target a known effector *CbNip1* of *C. beticola* (Ebert *et al*., 2021). Seedlings of BETA176 were infiltrated with BSCTV-*CbNip1* or BSCTV-*GFP*. At 40 dpi, we inoculated upper leaves with *C. beticola* by spraying method (Ebert *et al*., 2021). We observed highly reduced symptom formation for BSCTV-*CbNip1* infected plants compared to the control plant at 10 dpi (Figure 1j). In consistence, qPCR results showed significantly reduced fungal biomass in BSCTV-*CbNip1* infected plants compared to BSCTV-*GFP* infected plants (Figure 1k). Moreover, the relative expression levels of *CbNip1* in BSCTV-*CbNip1* treated plants were significantly down-regulated compared to those of control (Figure 1l), indicating that *CbNip1* was efficiently silenced by the cross-kingdom small RNAs. These results suggest that BSCTV VIGS system represents a platform to study sugar beet-pathogen interactions.

In summary, we have established an easy and efficiency BSCTV-based VIGS vector in sugar beet. This system will be an attractive tool for functional genomic studies for sugar beet and also possess the potential for a wide range of plant species.

## Materials and Methods

### Construction of BSCTV infectious clone and VIGS vectors

To construct the pCB-BSCTV plasmid, 1.0-copy and 0.2-copy fragments of the BSCTV genome were amplified by specific primer pairs BSCTV-1.0-F/R and BSCTV-0.2-F/R, respectively (Table S2). These two fragments were then ligated into the linearized pCB301 vector by homologous recombination. After transformation into *E. coli* DH5α, positive clones were screened by PCR, and the pCB-BSCTV plasmids were subsequently extracted from the cultured positive *E. coli*.

To engineer BSCTV into a VIGS vector, the pCB-BSCTV plasmid was linearized at a site one nucleotide downstream of the C1 protein, and 300-bp target sequence fragments were inserted at this position via homologous recombination. All primer pairs used in this study were listed in Table S2. Following *E. coli* transformation and PCR screening, positive plasmids of VIGS vectors targeting different genes were obtained.

### Agroinfiltration of *B. vulgaris*

To establish infection in sugar beet, the positive plasmids of each clone were transformed into *A. tumefaciens* GV3103. Positive clones were screened by PCR and cultured in LB medium containing kanamycin (50 mg/L) and rifampicin (25 mg/L) at 28°C with shaking at 220 rpm for 16 h. Cultured cells were harvested by centrifugation at 5,000 rpm for 5 min, resuspended in infiltration buffer [10mM MES (pH 5.6), 10 mM MgCl_2_, and 150 μM acetosyringone]. The bacterial suspension was then adjusted to an OD_600_ of 0.8 and incubated for 3 h prior to agroinfiltration.

### Plant Materials

All the test sugar beet varieties were cultivated in a growth chamber at 25°C under a 16 h/8 h light/dark cycle. Plants were used for subsequent experiments when the first pair of true leaves had fully expanded. Cells of *A. tumefaciens* containing BSCTV infectious clone or VIGS vectors were infiltrated into the sugar beet test plants at the first pair of true leaves. After several weeks of infection, systemic leaf tissues were collected from the plants for RT-qPCR analysis.

### Real-time quantitative RT-PCR (RT-qPCR)

Total RNA was extracted from systemic leaf tissues of BSCTV-infected plants and treated with recombinant DNase I (Takara) to remove DNA contamination. Subsequently, the DNA-free total RNA was reverse-transcribed into cDNA using M-MLV reverse transcriptase (Promega) with an oligo(dT) primer. Real-time qPCR reactions were performed using the 2×M5 HiPer Realtime PCR Super mix (Mei5bio) on QuantStudio1 Real-Time PCR System (Thermo Fisher).

## Acknowledgements

We thank Professors Jialin Yu, Dawei Li, Xian-Bing Wang, and Yongliang Zhang (China Agricultural University, Beijing, China) for their helpful comments on the research. We also thank Professor Xiaorong Tao (Nanjing Agricultural University, Nanjing, China) for providing the binary vector pCB301-2×35S-MCS-HDVRZ-NOS, Professor Huizhong Zhang (Inner Mongolia Academy of Agricultural & Animal Husbandry Sciences, Hohhot, China) for providing sugar beet varieties. This work was supported by the National Natural Science Foundation of China (32270165) and in part by China Agricultural Industry Technology System (Grant No. CARS-170304).

## Author contributions

Y.W., C.H., Z.Z. and Z.N. designed the experiments. Z.N., Z.G., X.Q., Q.L. and M.Z. conducted the experiments and analyzed the data. Z.N. and Y.W. wrote the manuscript.

## References

Ebert, M.K., Rangel, L.I., Spanner, R.E., Taliadoros, D., Wang, X., Friesen, T.L., de Jonge, R., Neubauer, J.D., Secor, G.A., Thomma, B.P., et al. (2021). Identification and characterization of cercospora beticola necrosis-inducing effector CbNip1. Mol. Plant Pathol. 22, 301–316.

FAO (2023). Crops and livestock products, 2023. Available at: https://www.fao.org/faostat/en/#data/QCL (accessed 9 July 2025)

Hatlestad, G.J., Akhavan, N.A., Sunnadeniya, R.M., Elam, L., Cargile, S., Hembd, A., Gonzalez, A., McGrath, J.M., and Lloyd, A.M. (2015). The beet Y locus encodes an anthocyanin MYB-like protein that activates the betalain red pigment pathway. Nat. Genet. 47, 92–96.

Liu Y, Schiff M, Dinesh-Kumar SP. (2002). Virus-induced gene silencing in tomato. Plant J. 31, 777–86.

Strausbaugh, C.A., Wintermantel, W.M., Gillen, A.M., and Eujayl, I.A. (2008). Curly top survey in the western United States. Phytopathology 98, 1212–1217.

Voorburg, C.M., Bai, Y., and Kormelink, R. (2021). Small RNA profiling of susceptible and resistant Ty-1 encoding tomato plants upon tomato yellow leaf curl virus infection. Front. Plant Sci. 12, 757165.

Zhang, Z., Chen, H., Huang, X., Xia, R., Zhao, Q., Lai, J., Teng, K., Li, Y., Liang, L., Du, Q., et al. (2011). BSCTV C2 attenuates the degradation of SAMDC1 to suppress DNA methylation-mediated gene silencing in arabidopsis. Plant Cell 23, 273–288.

