## Supplementary Material for "An efficient beet severe curly top virus-based VIGS vector in *Beta vulgaris*"

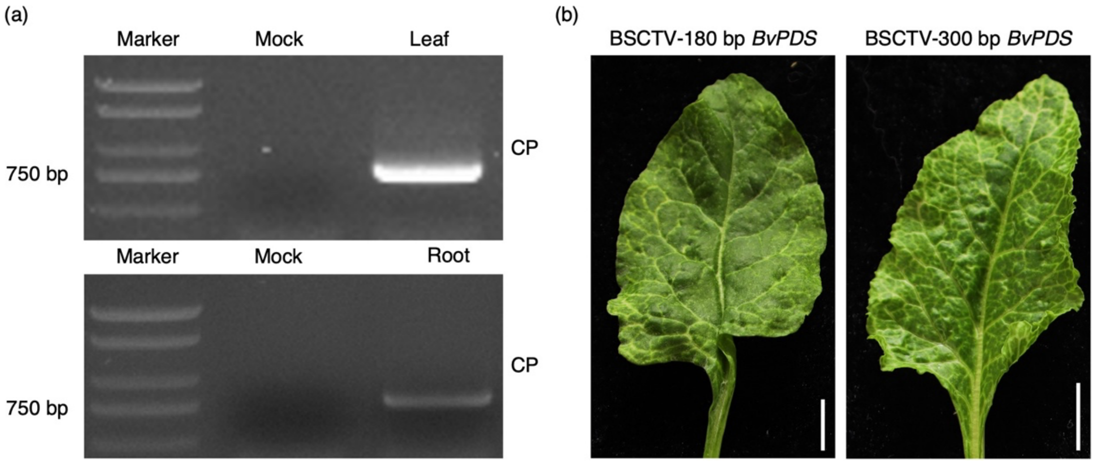


Figure S1: The BSCTV-based VIGS system in sugar beet. (a) The PCR detection of BSCTV-CP in the leaves and roots of sugar beet after agroinfiltration with BSCTV-wt. (b) The bleaching phenotype was observed on sugar beet leaves after agroinfiltration with VIGS vectors containing insertions of the 180 bp and 300 bp *BvPDS* sequences. Scale bars, 1 cm.

Table S1: The silencing efficiency of BSCTV-*BvPDS* in sugar beet BETA176.

| Number of plants for agroinfiltration | Number of silenced plants | Silencing efficiency |
|---|---|---|
| 55 | 45 | 81.82% |

Table S2: The primers used in this study.

| Primer | | Purpose |
|---|---|---|
| pCB301-F | TCGAATTTCCCCGATCGTTCAAACATTTGGCAATAAAGTTTCTTAAG | Construction of pCB301-BSCTV-wt |
| pCB301-R | CCTCTCCAAATGAAATGAACTTCCTTATATAGAGGAAGGGTCTTGC | |
| BSCTV-1.0-F | GGAAGTTCATTTCATTTGGAGAGGATTGAATCGGGCTCTCTTCA | |
| BSCTV-1.0-R | ATGCCTTTTTACAAAAAAGCCAAAAATTTTTTCCTTAC | |
| BSCTV-0.2-F | GGAAGTTCATTTCATTTGGAGAGGATTGAATCGGGCTCTCTTCA | |
| BSCTV-0.2-R | GAACGATCGGGGAAATTCGACTAGAAAGGCTGGATAATTGTCTG | |
| BSCTV-CP-F | ATGAGGAAATATACAAGAAATACGTATACAATGTCC | BSCTV-CP detection |
| BSCTV-CP-R | TTAATAAAAATAACATCTACAATTGCCAAACAAAGTG | |
| BSCTV-VIGS-F | TGTCATTTCACAATTCATCATGAATGTAATCAGGGATT | Linearization for fragment insertion |
| BSCTV-VIGS-R | CTTACAAGGAAGTTTGATCTTGCGAGGACG | |
| VIGS-BvPDS-F | AGATCAAACTTCCTTGTAAGTCGATCCATGCTGGAATTGG | Insertion fragments (Through homologous recombination) |
| VIGS-BvPDS-R | TGATGAATTGTGAAATGACAAGAACAGCACCCTCCATAGA | |
| VIGS-GFP-F | AGATCAAACTTCCTTGTAAGTGGAGTTGTCCCAATTCTTG | |
| VIGS-GFP-R | TGATGAATTGTGAAATGACACGTGTCTTGTAGTTCCCGTC | |
| VIGS-BvMYB1-F | AGATCAAACTTCCTTGTAAGGTTGTAAAGGGTTCATGGTCAGAT | |
| VIGS-BvMYB1-R | TGATGAATTGTGAAATGACAATGAGTCATAGTGTTTGTAGCTGG | |
| VIGS-CbNip1-F | AGATCAAACTTCCTTGTAAGATGAAGACCACCATCATCCT | |
| VIGS-CbNip1-R | TGATGAATTGTGAAATGACACTAGCAAGTTCCACGGTAAC | |

| | | |
|---|---|---|
| VIGS-test-F | GCCCTTAGGTCCTGGACATTACAAAATT | Insertion site test |
| VIGS-test-R | ATTTGCGGAGGTTGTGGTTGAATCT | |
| Bvactin-qF | ACATCAGCCGAACGGGAAAT | qPCR |
| Bvactin-qR | CCGATCAATGAGGGCTGGAA | |
| BvPDS-qF | TGCTTATGTCTCTGCGGCTC | |
| BvPDS-qR | AGCGAACTCCTGCTGAAGAG | |
| BvMYB1-qF | CGAAAACATTCCCGCACCAG | |
| BvMYB1-qR | TGCCCCAAATTATCCTGCTCA | |
| Cbactin-qF | ACATGGCTGGTCGTGATTTG | |
| Cbactin-qR | TGTCCGTCAGGAAGCTCGTA | |
| CbNip1-qF | CGCATTGATGGCGATGCTTT | |
| CbNip1-qR | TCCTTCAAACGTACTCCTCCG | |
